# Structural and Functional Characterization of the Alanine Racemase from *Streptomyces coelicolor* A3(2)

**DOI:** 10.1101/089938

**Authors:** Raffaella Tassoni, L.T. van der Aart, M. Ubbink, G. P. Van Wezel, N. S. Pannu

**Affiliations:** Leiden Institute of Chemistry, Leiden University, Gorlaeus Laboratories, Eisteinweg 55, 2333 CC Leiden, The Netherlands; Microbial Biotechnology and Health, Institute of Biology Leiden, Leiden University, Sylviusweg 72, 2333 BE Leiden, The Netherlands

**Keywords:** Alanine racemase, Peptidoglycan, Actinobacteria, Antibiotic resistance, X-ray crystallography

## Abstract

The conversion of L-alanine (L-Ala) into D-alanine (D-Ala) in bacteria is performed by pyridoxal phosphate-dependent enzymes called alanine racemases. D-Ala is an essential component of the bacterial peptidoglycan and hence required for survival. The Gram-positive bacterium *Streptomyces coelicolor* has at least one alanine racemase encoded by *alr*. Here, we describe an *alr* deletion mutant of S. *coelicolor* which depends on D-Ala for growth and shows increased sensitivity to the antibiotic D-cycloserine (DCS). The crystal structure of the alanine racemase (Alr) was solved with and without the inhibitors DCS or propionate, at 1.64 Å and 1.51 Å resolution, respectively. The crystal structures revealed that Alr is a homodimer with residues from both monomers contributing to the active site. The dimeric state of the enzyme in solution was confirmed by gel filtration chromatography, with and without L-Ala or D-cycloserine. Specificity of the enzyme was 66 +/− 3 U mg^−1^ for the racemization of L-to D-Ala, and 104 +/− 7 U mg^−1^ for the opposite direction. Comparison of Alr from S. *coelicolor* with orthologous enzymes from other bacteria, including the closely related D-cycloserine-resistant Alr from S. *lavendulae*, strongly suggests that structural features such as the hinge angle or the surface area between the monomers do not contribute to D-cycloserine resistance, and the molecular basis for resistance therefore remains elusive.

## INTRODUCTION

All canonical, proteinogenic amino acids, with the exception of glycine, have a stereocenter at the Cα and can exist either as the L-or D-enantiomer. While in the past it was generally accepted that only L-amino acids had a role in living organisms, studies revealed a variety of roles for free D-amino acids, for example in the regulation of bacterial spore germination and peptidoglycan structure [1]. Peptidoglycan is essential for cell shape and osmotic regulation and its biosynthesis is dependent on the availability of the less naturally abundant D-alanine [2, 3]. D-alanine (D-Ala) is generated through racemization of the abundant L-enantiomer (L-Ala) by the enzyme alanine racemase [4].

Alanine racemases (Alr; E.C. 5.1.1.1) are conserved bacterial enzymes that belong to the Fold Type III of pyridoxal phosphate (PLP)-dependent enzymes. Crystallographic studies of Alr’s from different bacterial species revealed a shared, conserved fold consisting of an eight-stranded α/β barrel at the *N*-terminal domain and a second, C-terminal domain mainly consisting of β-sheets. The PLP cofactor is bound to a very conserved lysine residue at the *N*-terminal side of the last helix of the barrel. In all studies, Alr’s crystals consist of homo-dimers, with residues from both monomers participating in the formation of the active site. Nevertheless, in-solution studies indicated an equilibrium between monomeric and homo-dimeric states in solution, with the homo-dimer being the catalytically active form [5, 6].

*Streptomyces coelicolor* is the most widely studied member of the Streptomycetes, which are Gram-positive bacteria with a multicellular mycelial life style that reproduce via sporulation [7]. They are of great importance for medicine and biotechnology, accounting for over half of all antibiotics, as well as for many anticancer agents and immunosuppressants available on the market [8, 9]. Here we describe the heterologous expression, purification and crystal structure of *S. coelicolor* Alr, encoded by *alr* (SCO4745). An *alr* null mutant of *S. coelicolor* depends on exogenous D-Ala for growth. The purified enzyme catalyzed the racemization of L-Ala to D-Ala *in vitro* and was shown to be a dimer from gel filtration, both in the absence and presence of L-Ala or the inhibitor D-cycloserine (DCS). Furthermore, the crystal structure of the enzyme has been solved both in the absence and in the presence of the inhibitors DCS and propionate. The comparison of the Alr enzymes from *S. coelicolor* and its close relative the DCS-resistant *Streptomyces lavendulae* [10] questions the structural role of Alr in DCS resistance.

## MATERIALS AND METHODS

### Bacterial strains and culturing conditions

*Escherichia coli* strains JM109 [11] and ET12567 [12] were used for routine cloning procedures and for extracting non-methylated DNA, respectively. *E. coli* BL21 Star (DE3)pLysS was used for protein production. Cells of *E. coli* were incubated in lysogenic broth (LB) at 37°C. *Streptomyces coelicolor* A3(2) M145 was obtaime from the John Innes cenre strin collection and was the parent of all mutants in this work. All media and routine *Streptomyces* techniques have been described in the *Streptomyces* laboratory manual [12]. Soy, flour and mannitol (SFM) agar plates were used for propagating *S. coelicolor* strains and for preparing spore suspensions. Solid minimal medium (MM) agar plates supplied with 1% (w/v) mannitol was used for phenotypic characterization. The MIC for D-cycloserine was measured in triplicate in 96 well plates containing solid MM, D-alanine and D-cycloserine. Growth was assayed after two days using a resazurin-assay [13].

### Gene disruption and genetic complementation

*S. coelicolor alr* deletion mutants were constructed as previously described [14]. The −1238/−3 and +1167/+2378 regions relative to the start of *alr* were amplified by PCR using primer alr_LF and alr_LR, and alr_RF and alr_RR (Table S2) as described [15]. The left and right flanks were cloned into the unstable, multi-copy vector pWHM3 (Table S1) [16], to allow for efficient gene disruption [17]. The apramycin resistance cassette *aac(3)IV* flanked by *loxP* sites was cloned into the engineered XbaI site to create knock-out construct pGWS1151. Media were supplemented with 1 mM D-Ala to allow growth of *alr* mutants. The correct recombination event in the mutant was confirmed by PCR. For genetic complementation, the −575/+1197 region (numbering relative to the *alr* translational start codon) encompassing the promoter and coding region of SCO4745 was amplified from the *S. coelicolor* M145 chromosome using primer 4745FW-575 and 4745RV+1197 (Table S2) and cloned into pHJL401 [18]. pHJL401 is a low-copy number shuttle vector that is very well suited for genetic complementation experiments [19].

### Cloning, expression and purification of His_6_-tagged Alr

The *alr* gene (SCO4745) was PCR-amplified from genomic DNA of *Streptomyces coelicolor* using primers alr-FW and alr-RV (Table S2), and cloned into pET-15b with a *N*-terminal His_6_ tag. The construct was introduced into *E. coli* BL21 Star (DE3)pLysS (Novagen). His_6_-tagged Alr was purified using a Ni-NTA column (GE Healthcare) as described [20]. Fractions containing Alr were desalted using a PD-10 column (GE Healthcare) to remove the imidazole and Alr purified further by gel-filtration using a Superose-12 column (GE Healthcare). The fractions were analyzed by SDS-PAGE and those containing pure Alr were pooled, concentrated and flash-frozen in liquid nitrogen. The purified protein was verified by tandem mass spectrometry analysis using an LTQ-Orbitrap (Thermo Fisher Scientific, Waltham, MA) as previously described [21].

### Enzymatic assay and kinetics

0.2 nM purified Alr was added to 10 mM L-Ala in 20 mM PBS buffer, pH 7.0 and incubated at room temperature (21°C) for 0, 1, 5, 15, 30, and 60 min. The reaction was stopped by heat inactivation for 3 min at 95°C. As a control, 0.2 nM heat-inactivated Alr was used. A 20-μL aliquot of the reaction mixture was added to a 2-mL well in a 24-well plate containing solid MM, inoculated with the *alr* mutant. 0, 10 μM, 50 μM 100 μM, 500 μM and 1 mM D- and L-Ala were added as controls.

The racemization activity of Alr was determined by quantifying the derivatization product of L-and D-Ala by HPLC, as previously described [22, 23]. For detailed methods see the Supplementary Material.

### Crystallization conditions and data collection

Purified Alr was concentrated to 20 mg mL^−1^ and crystallization conditions were screened by sitting-drop vapor-diffusion using the JCSG+ and PACT *premier* (Molecular Dimensions) screens at 20°C with 500-nL drops. The reservoir (75 μL) was pipetted by a Genesis RS200 robot (Tecan). Drops were made by an Oryx6 robot (Douglas Instruments). After 1 day, crystals grew in condition number 2-38 of PACT *premier*, which consisted of 0.1 M BIS-TRIS propane, pH 8.5, 0.2 M NaBr, 20% (w/v) PEG3350. Bigger crystals were grown in 1-μL drops. Prior to flash-cooling in liquid nitrogen, crystals were soaked in a cryosolution consisting of mother liquor, 10% glycerol and 10, 5, or 1 mM inhibitor to obtain the DCS-and propionate-bound structures.

X-ray data collection was performed at the ESRF (Grenoble, France) on beamline ID23-1 [24] using a PIXEL, Pilatus_6M_F X-ray detector. A total of 1127 frames were collected for the native Alr, with an oscillation of 0.15°, an exposure time of 0.378 s, total 426.006 s. The data set was processed by XDS [24] and scaled by AIMLESS [25] to a resolution of 2.8 Å. For the DCS-bound Alr 960 frames were collected, with an oscillation range of 0.15°, an exposure time of 0.071 s per image and a total time of 68.16 s. For the propionate-bound structure, 1240 images were collected, with an oscillation degree of 0.1°, an exposure time per image of 0.037 s, and a total time of 45.88 s. Both inhibitor-bound data sets were auto-processed by the EDNA Autoprocessing package in the mxCuBE [26] to a resolution of 1.64Å and 1.51 Å for the DCS-bound and propionate-bound Alr, respectively. The structures were solved by molecular replacement with *MOLREP* [27] using 1VFH as a search model from the *CCP4* suite [28] and iteratively refined with REFMAC [29]. Manual model building was done using *Coot* [30].

## RESULTS AND DISCUSSION

### Creation of an *alr* null mutant of *S. coelicolor*

To study the role of *alr* in amino acid metabolism and in morphogenesis of *S. coelicolor*, a deletion mutant was created by removal of the entire coding region (see Materials & Methods). In line with its expected role in biosynthesis of the essential D-amino acid D-Ala, *alr* null mutants depended on exogenously added D-Ala (50 μM) for growth (Fig 1a). Reintroducing a copy of the *alr* gene via plasmid pGWS1151 complemented the mutant phenotype, allowing growth in the absence of added D-Ala (Fig 1a). These data show that *alr* encodes alanine racemase in *S. coelicolor*, and that it most likely is the only gene encoding this enzyme.

**Figure 1.**
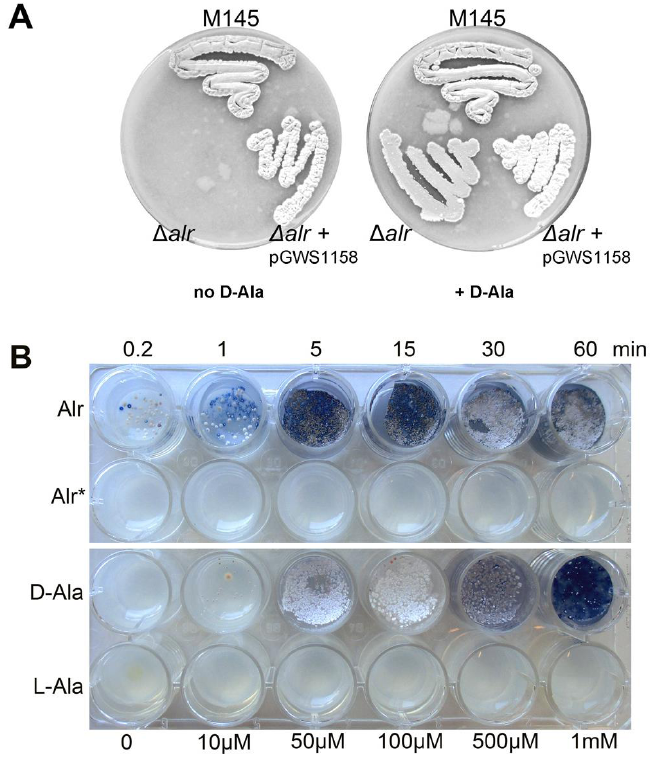
SCO4547 encodes alanine racemase Alr. (A) The *alr* null mutant of *S. coelicolor* requires supplemented D-Ala for growth. This phenotype is rescued by the addition of 1 mM D-Ala to the media or by the introduction of plasmid pGWS1151, which expresses Alr. Strains were grown for 5 days on SFM agar plates. (B) Rescue of the *S. coelicolor alr* null mutant with D-Ala produced *in vitro* by Alr. Alr (0.2 nM) was incubated with 10 mM L-Ala for 10 sec or 1, 5, 15, 30 or 60 min, followed by heat inactivation. The reaction mixture was then added to MM agar in 24-well plates and growth of the *alr* null mutant assessed. As a control, the same experiment was done with heat inactivated Alr (Alr*). Note the restoration of growth to the *alr* mutant after addition of the reaction mixture with active Alr, but not with heat-inactivated Alr. The bottom rows show control experiments with added D-Ala or L-Ala (ranging between 0-1 mM).

### Alr is the only alanine racemase in *S. coelicolor*

The Alr from *S. coelicolor* (Alr_Sco_) shares 74.9% aa sequence identity with the Alr from the DCS-producing *Streptomyces lavendulae* (Alr_Sla_) [10] and 37.9% with Alr from *Geobacillus stearothermophilus* (Alr_Gst_) [31] (Fig. 2). To analyze the activity and kinetics of the enzyme, recombinant His_6_-tagged Alr was expressed in *E. coli* BL21 Star cells and purified to homogenity (see Materials and Methods section). Around 45 mg of pure Alr was obtained from 1 L of bacterial culture and Alr was identified using mass spectrometry (Table S4).

**Figure 2.**
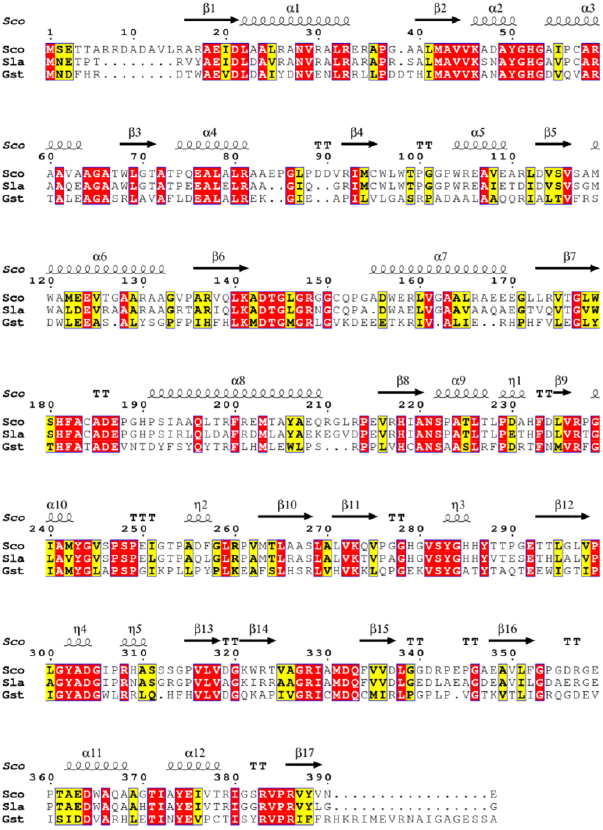
Multiple aminoacid sequence alignment of Alr from *S. coelicolor* (Sco), S. *lavendulae* (Sla; PDB 1VFT) and *G. stearothermophilus* (Gst; PDB 1XQK). The sequence alignment showing structural elements of Alr_Sco_ was generated with ESPript3.0. α-helices are shown as large coils labeled a, 3_10_-helices are shown as small coils labeled ŋ, β-strands are shown as arrows labeled β and β-turns are labeled TT. Identical residues are shown on a red background, conserved residues are shown in red and conserved regions are shown in boxes.

To establish if Alr_Sco_ converts L-Ala into D-Ala, we designed a combined *in vitro* and *in vivo* assay, whereby 10 mM L-Ala was incubated with 0.2 nM purified Alr-His_6_ for 10 sec to 60 min, after which the enzyme was heat-inactivated and the mixture was added to minimal media (MM) agar plates, followed by plating of 10^6^ spores of the *alr* null mutant and incubation of 5 days at 30°C (Fig. 1b). If Alr converts L-Ala into D-Ala, the reaction should generate sufficient D-Ala to allow restoration of growth to *alr* null mutants. Indeed, sufficient D-Ala was produced to allow growth of the *alr* null mutant, whereby biomass accumulaiton was proportional to the incubation time, while a control experiment with extracts from heat-inactivated Alr did not give any growth (Fig. 1b). Addition of D-Ala also restored growth of *alr* mutants, while no growth was seen for cultures grown in the presence of added L-Ala. Taken together, this shows that no Alr activity is present in *S. coelicolor alr* mutants, and that Alr actively converts L-Ala into D-Ala *in vitro*.

The kinetic parameters of recombinant Alr_Sco_ were determined using both L- and D-Ala as substrates. The enzyme shows a K_m_ of 6.3 mM and 8.9 mM towards L-and D-Ala, respectively (Table S5). For comparison, kinetic parameters of Alr_Sla_ [10] and Alr_Gst_ are also shown in Table S5. To analyze possible multimerization of Alr_Sco_, analytical gel filtration was used, which established an apparent molecular weight for the protein of 83 kDa in solution both in the absence and in the presence of L-Ala and DCS (Figure S2); this corresponds roughly to two Alr proteins (43.4 kDa per subunit). See Supplementary Data for more details. These data suggest that Alr_Sco_ forms a dimer in solution, both in the presence and in the absence of ligands.

### Structure of Alr from *Streptomyces coelicolor*

The crystal structure of ligand-free Alr_Sco_ was determined to a resolution of 2.8 Å (Table S6). The structure is similar to that of other prokaryotic Alr proteins [10, 31]. The asymmetric unit contains four protein molecules (A-D), which interact head-to-tail to form two dimers (dimer A-B and C-D, Fig. 3a). Each monomer has two structurally distinct domains. The *N*-terminal domain, comprising residues 1-259, has an eight-stranded α/β-barrel fold, typical of phosphate-binding proteins. The C-terminal domain, residues 260-391, contains mostly β-strands (Fig. 3b). 96% of the amino acids are in the preferred regions of the Ramachandran plot, 4% are in the allowed regions and no residues are in the unfavored areas. After refinement, the r.m.s. deviation between Cα’s of the two interacting molecules A and B is 0.0794 Å, and between C and D is 0.1431 Å.

**Figure 3.**
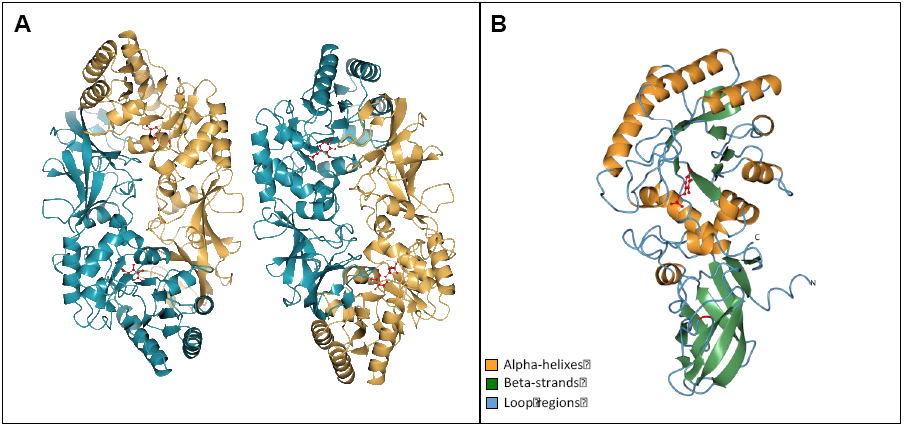
Ribbon representation of Alr from *S. coelicolor*. (A) The asymmetric unit contains two homo-dimers. The two chains within one dimer are colored in gold and dark cyan. (B) The *N*-terminal domain (1-259) is an α/β-barrel. The C-terminal domain (260-391) mostly comprises β-strands. The catalytic Tyr283 is highlighted in red.

There are two active sites per dimer, which are located at the interface between each α/β-barrel of one subunit and the *C*-terminal domain of the other. The catalytic core (Fig. 4a) consists of the PLP cofactor, a Lys, and a Tyr, which is contributed by the other subunit. The PLP is bound through an internal aldimine bond to the amino group of Lys46, located at the C-terminal side of the first β-strand of the α/β-barrel. The side chain of the catalytic Lys46 points out of the α/β-barrel, towards the C-terminal domain of the interacting subunit, and in particular, towards Tyr283’. The phosphate group of the PLP is stabilized by hydrogen bonds with the side chains of Tyr50, Ser222 and Tyr374, and with the backbone of Gly239, Ser222 and IIe240. The pyridine ring of the PLP is stabilized by a hydrogen bond between the N-1 of the cofactor and Nε of Arg237. The C2A of the PLP also interacts with oxygen Q1 of the carboxylated Lys141. All residues stabilizing the PLP cofactor (Tyr50, Ser222, Gly239, Ile240, Arg237, Tyr374) are conserved among Alr’s [31]. However, the Alr_Sco_ lacks one important hydrogen bond between Arg148 and the phenolic oxygen of the PLP molecule, as already observed for the Alr_Sla_ [32] [33]. Among the residues involved in PLP-stabilization, Tyr374, which corresponds to Tyr354 in the well-studied Alr_Gst_, is particularly interesting. In fact, while always regarded as a PLP-stabilizer, Tyr354 was shown to be actually anchored by the hydrogen bond with the PLP so to ensure substrate specificity of the Alr [34].

**Figure 4.**
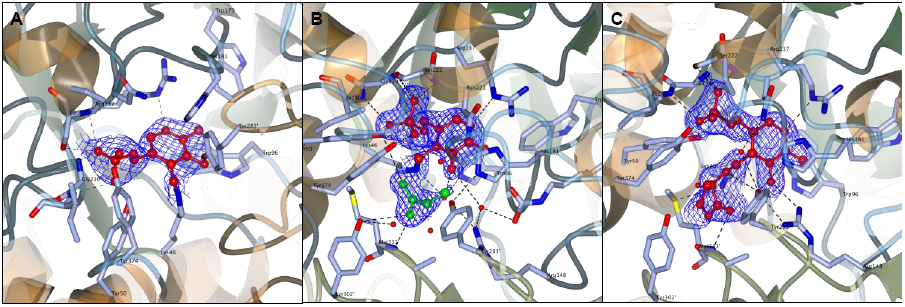
Active site of Alr without ligands (A) and in complex with propionate (B) and D-cycloserine (C). The electron density 2F_0_-F_c_ map is shown for the PLP and the two inhibitors. The residues in the active site which are contributed from the monomer not bound to the PLP are indicated as primed numbers. Dotted lines indicate interatomic distances shorter than 3Å.

### Alr crystal structures bound to propionate and DCS

The structure of Alr_Sco_ was solved in the presence of the two inhibitors propionate [35] and DCS [10] to a resolution of 1.51 Å and 1.64 Å, respectively. In Alr_Sco_, the carboxylic group of propionate binds in the same orientation in all four active sites present in the crystal unit. One of the carboxylic oxygens of propionate forms hydrogen bonds with the amino group of Met331’, the phenolic oxygen and CE2 of Tyr283’ and a water molecule, which bridges the substrate to several other water molecules in the active site. The second carboxylic oxygen of propionate interacts with Nζ of Lys46, C4A of PLP, the phenolic oxygen of Tyr283’ and a water molecule. The electron density around C3 of propionate shows that this atom can have several orientations in the active site. In fact, it can either interact with O4 of the phosphate of PLP, the phenolic oxygen of Tyr283’ and a water molecule, or with the phenolic oxygen of Tyr50, the CE of Met331’, Nζ of Lys46 and C4A of PLP (Fig. 4b). Both in Alr_Sco_ and Alr_Gst_ the hydrogen-bonding network stabilizing the propionate is very conserved. The propionate-bound form of the Alr can be considered a mimic of the Michaelis complex formed between the enzyme and its natural substrate L-Ala [35].

DCS is an antibiotic produced by S. *lavendulae* and *Streptomyces garyphalus* and covalently inhibits Alr [10, 36]. The structure of the Alr_Sco_ in complex with DCS shows that the amino group of the inhibitor replaces the Lys46 in forming a covalent bond with the PLP C4 (Fig. 4c). The nitrogen and oxygen atoms in the isoxazole ring form hydrogen bonds with Tyr302’, Met331’ and a water molecule. The hydroxyl group of the DCS ring also interacts with Met331’ and Arg148 (Fig. 4c). The catalytic Tyr283’ is at hydrogen-bond distance from the amino group of DCS. DCS molecules superpose well in three of the active sites of the crystal (A, B and C). However, the DCS-PMP adduct in the fourth site showed a shift of the DCS compared to the other sites, while PLP still superposed well. Comparison of the hydrogen bonds in the proximity of the active site in the free and in the DCS-bound form of Alr, revealed a role for Arg148 in substrate stabilization. In fact, while the side chain of Arg148 is hydrogen-bonded to Gln333 in free Alr, it shifted towards the hydroxyl moiety of the DCS upon binding of the inhibitor. Gln333 was stabilized by interaction with carboxylated Lys141. The rearrangement of the hydrogen bonds involving Arg148, Gln333 and Lys141 does not occur in the propionate-bound structure, suggesting that Arg148 is involved in stabilization of the carboxylic group of the substrate.

### Alr and DCS resistance

Alr_Sco_ shows high sequence and structural similarity to Alr_Sla_. Based on crystallographic studies, Alr_Sla_ was proposed as one determinant of DCS resistance in the producer strain [10]. However, detailed structural comparison of different Alr proteins based on interatomic distances between active site residues, dimerization areas and hinge angles (see Supplementary Data), failed to establish a clear correlation between structural features and DCS resistance. It therefore remains unclear what the role of Alr is in DCS resistance in the producer strain.

In conclusion, our work shows that SCO4745 encodes the only Alr in *S. coelicolor* A3(2) and is essential for growth. Our structural studies on Alr_Sco_ and comparison to Alr_Sla_ suggest that there are more factors contributing to DCS resistance in bacteria, which relates well to recent studies on DCS-producing streptomycetes [37] and DCS-resistant strains of *M. tuberculosis* [38–41]. The availability of a *S. coelicolor alr* null mutant and the crystal structure of the Alr from the same bacterium offer a new, valuable model for the study of DCS resistanc

